# Mechanical force locally damages, remodels and stabilizes the lattice of spindle microtubules

**DOI:** 10.1101/2025.06.05.657915

**Authors:** Caleb J. Rux, Megan K. Chong, Valerie Myers, Nathan H. Cho, Sophie Dumont

## Abstract

To segregate chromosomes at cell division, the spindle must maintain its structure under force. How it does so remains poorly understood. To address this question, we use microneedle manipulation to apply local force to spindle microtubule bundles, kinetochore-fibers (k-fibers), inside mammalian cells. We show that local load directly fractures k-fibers, and that newly created plus-ends often have arrested dynamics, resisting depolymerization. Force alone, without fracture, is sufficient for spindle microtubule stabilization, as revealed by laser ablating k-fibers under local needle force. Doublecortin, which binds a compacted microtubule lattice, is lost around the force application site, suggesting local force-induced structural remodeling. In turn, EB1, which recognizes GTP-tubulin, is locally enriched at stabilization sites, both before and after force-induced fracture. Together, our findings support a model where force-induced damage leads to local spindle microtubule lattice remodeling and stabilization, which we propose reinforces the spindle where it experiences critical loads.

## Introduction

Cell division is essential to life. Each time a cell divides, it builds the spindle, a cellular machine composed of microtubules, motors and crosslinkers. The spindle is responsible for faithful chromosome segregation, a process that happens over about an hour in mammals. Throughout this time, it must generate and respond to mechanical force to build and maintain its structure, ensure correct chromosome attachments and ultimately segregate chromosomes^1–4^. How the spindle maintains its structure in the face of forces it and its environment generate remains an open question.

We now have a nearly complete list of mammalian spindle components and understand mechanisms of force generation for many. We know from *in vitro* work how motors^5,6^ and microtubule dynamics^7–12^ generate force, and from *in vivo* work how their force generation play a role in the spindle^8,13–17^. *In vitro* work has revealed how force impacts motor^6,18,19^ and nonmotor^20,21^ proteins as well as microtubule dynamics^22,23^. However, our understanding of how spindle components respond to force in the spindle environment – and how the spindle ultimately responds to force – is more rudimentary. In part, this is because while we have many molecular tools to perturb force generators *in vivo*, exerting controlled, local force to probe how the mammalian spindle responds to force remains more challenging.

Key to chromosome movement inside the spindle^11^, the end dynamics of microtubules respond to force. *In vitro*, mechanical force regulates microtubule plus-end polymerization and depolymerization rates and the transitions between these states^22–24^. In vivo, force dictates the plus-and minus-end dynamics of mammalian k-fibers, a bundle of 10-30 kinetochore-bound microtubules^25–30^. These microtubule bundles are the longest (with many microtubules reaching both kinetochores and poles^25^) and longest-lived^31^ in the spindle; as such, they are poised to play a unique mechanical role in the spindle. How microtubule dynamics and biochemistry respond to force within a k-fiber remains a frontier, especially away from microtubule ends. Mammalian k-fibers cannot currently be reconstituted *in vitro*.

At interphase and *in vitro*, individual microtubules are not only dynamic at their ends but also along their lattice^32,33^. Mechanical force *in vitro,* motor proteins walking along microtubules, and laser damage of the lattice all encourage the exchange of GDP-tubulin for fresh GTP-tubulin dimers in individual microtubules, far from microtubule ends^32–36^. Local biochemical remodeling and stabilization of the microtubule lattice can accompany such tubulin exchange *in vitro* and *in vivo* at interphase^32–37^. Given that k-fiber spindle microtubules are often under load, force could also remodel and stabilize the lattice of k-fiber microtubules in the spindle^4^. Indeed, microneedle manipulations of k-fibers over minutes-long timescales suggested their lattice stability may differ under load^30,38^. Whether and how local force does so, over what spatial and temporal scales it acts, and its impact on lattice biochemistry and dynamics, remain open questions. More broadly, whether the bundling of microtubules, such as those in k-fibers, impacts such lattice remodeling is also an open question.

Here we show that mechanical force locally damages the lattice of spindle microtubules, leading to local microtubule remodeling and stabilization. To do so, we use a combination of microneedle manipulation, laser ablation, and live-cell imaging to physically interrogate spindle microtubules in PtK2 cells. We show that spindle microtubule bundles fracture along their length under local load and that their new plus-ends often display arrested dynamics, neither shrinking nor growing appreciably. We demonstrate that force alone, independent of and before fracture, can stabilize spindle microtubules and change their lattice biochemistry. Spindle microtubules exhibit local doublecortin loss at the site of force application and local EB1 enrichment at stabilization sites before and after fracture. Thus, force locally remodels spindle microtubule lattice structure, biochemistry and dynamics – stabilizing microtubules. We propose that this force induced damage, repair and stabilization reinforces the spindle where it is under critical loads, helping maintain its structure in the face of mitotic and environment forces.

## Results

### Sustained local load directly fractures spindle microtubules and stabilizes new microtubule plus-ends

To probe how microtubules respond to local force in the mammalian spindle, we used microneedle manipulation in PtK2 cells (Figure 1a). Microneedle manipulation allows us to apply local loads to individual k-fibers with high temporal and spatial precision while preserving cell health^15,30,39^. In turn, rat kangaroo PtK2 cells offer a low chromosome number for a mammal, allowing us to reproducibly pull on a single k-fiber. Using a fluorescently coated glass microneedle, we pulled on an outer k-fiber in spindles of PtK2 GFP-tubulin cells^40^, pulling over 15 µm in 216 s (4 µm/min manipulation speed) (Figure 1b). As expected from our previous work^30^, only manipulated k-fibers appeared to on average lengthen under sustained force (Figures 1c-d, Supplementary Figure 1, Videos 1-2). About 60% of k-fibers fractured along their length under this sustained load (Figure 1e), and we never observed k-fiber detachment from spindle poles or kinetochores^30^. To verify microtubule fracture, rather than k-fiber movement out of plane or sliding apart of microtubules within the bundle, we imaged the full cell volume after removing the microneedle from the cell. Consistent with fracture, we typically saw mechanical uncoupling of the k-fiber stubs, i.e. both stubs taking dramatically different orientations after fracture events. Further supporting fracture, we could not detect microtubules between the two k-fiber stubs (Figure 1c-d).

**Figure 1:**
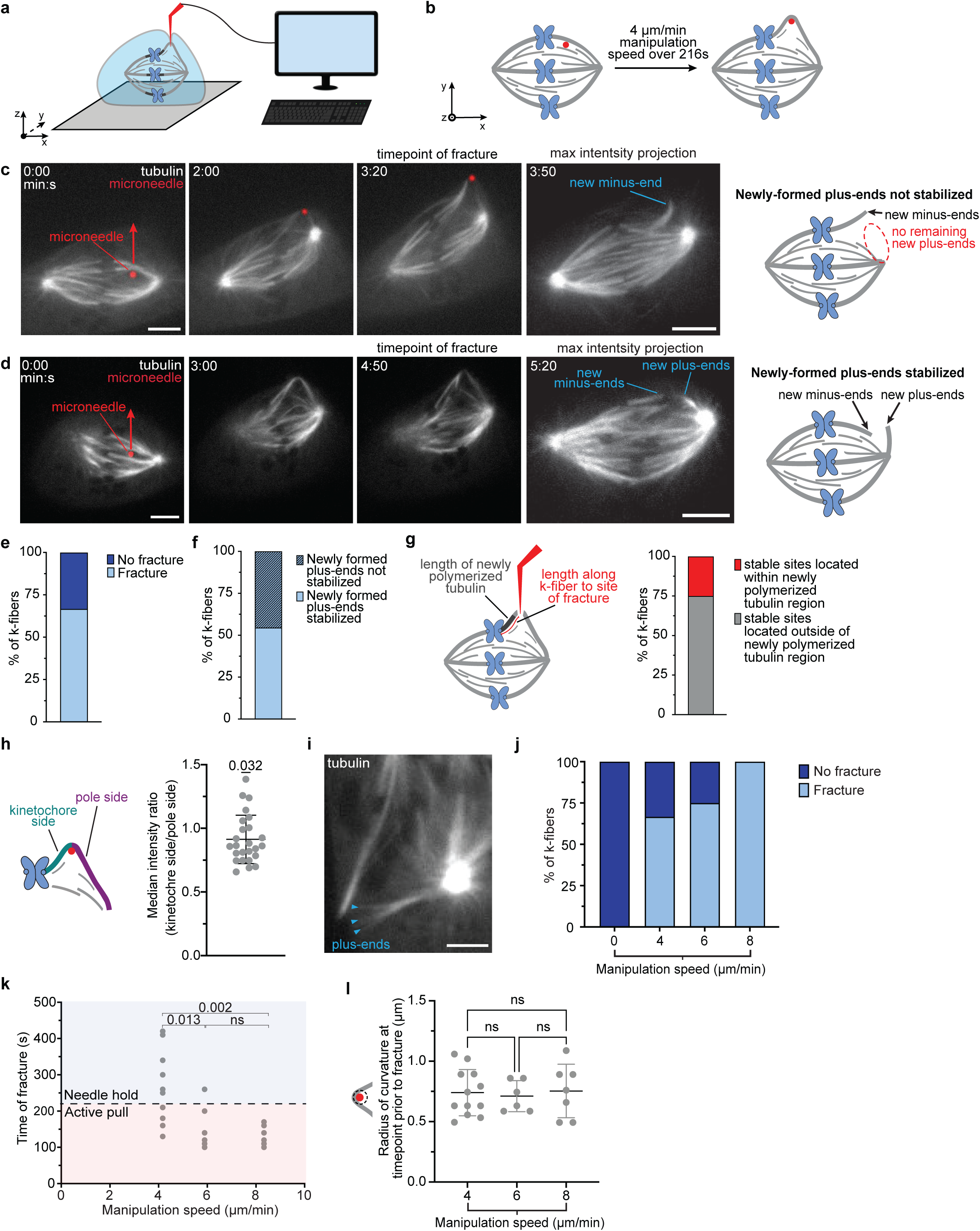
Sustained local load directly fractures spindle microtubules and stabilizes new microtubule plus-ends. **a)** Schematic of microneedle manipulation setup, with needle (red) movement in x, y and z being computer-controlled to generate reproducible perturbations on the mammalian spindle. **b)** Schematic depicting microneedle manipulation locally exerting force on a k-fiber in the metaphase spindle. **c & d)** A representative timelapse (single plane) of a k-fiber fracture and lack of stabilization (c), or subsequent stabilization (d), in PtK2 cells stably over-expressing GFP-tubulin (white) where the microneedle (red, AF647 labelled) traveled 15 µm (red arrow direction) over 216 s starting at time 0:00. Newly created plus-and minus-ends are indicated on the max intensity projection image (right). Scale bars = 5 µm. **e)** The percentage of total k-fibers that fractured when subject to microneedle manipulation over a distance of 15 µm over 216 s (n=18 k-fibers, N=7 experimental days). **f)** The percentage of fractured k-fibers that resulted in a newly formed stable plus-end (n=11 k-fibers, N=7 experimental days). **g)** The proportion of k-fiber fracture and stabilization events that occur within and outside of the newly polymerized tubulin region, as depicted in the adjacent spindle schematic (based on k-fiber length increase during manipulation) (n=25 k-fibers, N=9 individual experiments). **h)** The ratio of median k-fiber intensity between the kinetochore and the manipulation site and the pole and the manipulation site (pole side/kinetochore side) the timepoint preceding fracture (one sample t-test, n=25 k-fibers, N=9 individual experiments, mean tested for significant difference from 1). **i)** An image of splayed microtubule ends terminated at the same position along the k-fiber stub, consistent with a single fracture site and event. Scale bar = 3 µm. **j)** The percentage of total k-fibers that fractured at different loading rates. (n_0_=4, n_4_=18, n_6_=8, n_8_=7 k-fibers, N=9 experimental days). **k)** The time at which a fracture occurred (if it occurred) at different loading rates. (n_4_=12, n_6_=6, n_8_=7 k-fibers, N=9 experimental days, ordinary one-way ANOVA). **l)** K-fiber radius of curvature at the manipulation site one timepoint prior to fracture at different loading rates (n_4_=12, n_6_=6, n_8_=7 k-fibers, N=9 experimental days, ordinary one-way ANOVA).

About 50% of pole-side fractured k-fiber stubs (with new plus-ends) remained stable after load application (Figure 1f). These new plus-ends apparently resisted depolymerization for tens of seconds to minutes post fracture, as suggested by our previous work^30^. In contrast, acute severing of a k-fiber using laser ablation results in new plus-ends rapidly depolymerizing over seconds^41^, presumably due to the lack of a stabilizing GTP-tubulin “cap” region far from microtubule plus-ends at kinetochores. A simple model for the stability of newly created plus-ends is that newly polymerized GTP-tubulin, fresh from plus-end addition under load at kinetochores, moved to the fracture site as the k-fiber polymerized from its plus-end and stopped minus-end depolymerization under load^30^. If this were the case, the sites of the newly formed plus-ends would need to be within this newly polymerized region, measured by overall k-fiber growth. However, 75% of k-fibers did not grow enough to account for a GTP-tubulin region near the fracture site (Figure 1g). Thus, the stability of new spindle microtubule plus-ends is not simply due to newly incorporated GTP-tubulin at kinetochore-bound plus-ends reaching the future fracture site, suggesting stabilization is instead locally conferred.

In principle, spindle microtubules could fracture as a direct consequence of force or of k-fiber microtubules having a finite lifetime^31^ and not being replenished under manipulation. Several observations support the former model. First, k-fiber stubs often rapidly change plane upon fracture, suggesting an abrupt rather than gradual breakage event. Second, k-fibers exhibit only a small, but statistically significant reduction in intensity (∼10%) on the pole side compared to the kinetochore side of the manipulation site just prior to fracture (Figure 1h). If gradual microtubule loss caused fracture, for example because potential k-fiber microtubules cannot “turn the corner” around the needle, microtubule intensity on one k-fiber side should instead approach zero. Third, individual microtubule ends often splayed out from one or both stubs and splayed ends terminated at the same position along the stub (Figure 1i), suggesting a single k-fiber fracture site and event. Fourth, to more directly test these two models we subjected individual k-fibers to different manipulation speeds (0, 4, 6, and 8 µm/min) and thus loading rates (rate of change of force) over the same period (216 s) followed by maintaining the final needle position, maintaining force on the k-fiber, until the k-fiber either fractured or >7 mins elapsed from the start of manipulation; the 0 µm/min speed controlled for the presence of the needle, rather than its movement, causing fracture. Higher loading speeds over the same period result in higher magnitudes of force in a viscoelastic material^42^, like the spindle^43^. The likelihood of fracture increased (Figure 1j) and time to fracture decreased (Figure 1k) with loading rate, consistent with force directly causing fracture and not easily consistent with microtubule turnover causing apparent fracture. Fifth, k-fibers fractured at a similar radius of curvature regardless of loading rate (Figure 1l), consistent with a similar magnitude of force achieved prior to fracture across all conditions. Together, these findings indicate that local load directly fractures k-fiber spindle microtubules and that the newly formed plus-ends often appear to resist depolymerization.

### Sustained local load stabilizes spindle microtubules independent of fracture

If mechanical force directly leads to fracture (Figure 1), it could lead to lattice damage and stabilization^32,33^ at the future fracture site and new plus-ends (Figure 1d-i), before fracture. Alternatively, stabilization of the new plus-ends could occur only after they are created, and not before fracture. To determine if k-fiber lattice stabilization occurs prior to and independent of fracture, we sought to exert local force and query the state of the microtubule lattice without force-induced fracture. We used laser ablation to sever k-fibers to query the lattice state. When we laser ablated unmanipulated k-fibers (never contacted by the microneedle or contacted and never manipulated) in PtK2 GFP-tubulin cells, all rapidly depolymerized, as expected (Figure 2a-b, Video 3)^41^.

**Figure 2:**
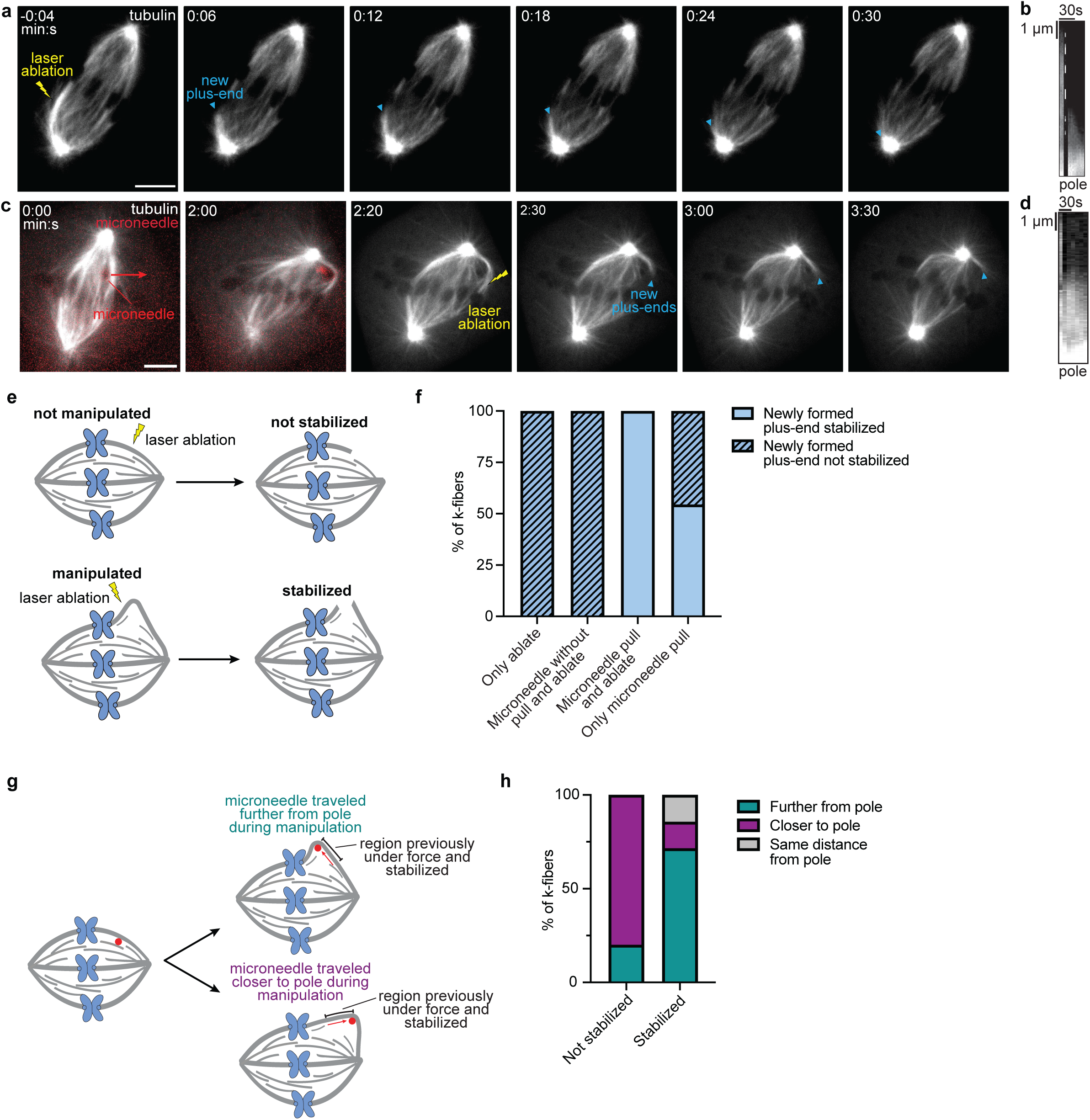
Sustained local load stabilizes spindle microtubules independent of fracture. **a)** Representative timelapse of an unperturbed metaphase k-fiber severed via laser ablation (at time 0:00, yellow), with plus-ends (blue arrowhead) rapidly depolymerizing to the spindle pole in PtK2 cells stably overexpressing GFP-tubulin (white). Scale bar = 5 µm. **b)** Kymograph of the ablated k-fiber, demonstrating rapid plus-end depolymerization. **c)** Representative timelapse of a metaphase k-fiber subject to microneedle manipulation (red arrow, starting at time 0:00, 4 μm/min pull speed, AF647 labelled) before being severed via laser ablation (yellow) on the kinetochore side of the manipulation site in PtK2 cells stably overexpressing GFP-tubulin (white). The newly formed plus-ends (blue arrowhead) demonstrate stabilization only seen when microneedle load is applied. Scale bar = 5 µm. **d)** Kymograph of the manipulated and ablated k-fiber, demonstrating a lack of depolymerization over tens of seconds. **e)** Schematic depicting the two possible outcomes of laser ablation without (top) and with (bottom) microneedle pulling: either new plus-ends are stabilized or they are not. **f)** The percentage of fractured k-fibers that resulted in newly formed stable plus-ends when severed via laser ablation (“Only ablate”), with the microneedle next to the k-fiber for >2 mins without pulling followed by laser ablation (“Microneedle without pull and ablate”), microneedle manipulation for ∼2 mins followed by microneedle removal and laser ablation (“Microneedle pull and ablate”), or 4 µm/min microneedle manipulation alone (“Only microneedle pull”). **g)** Schematic depicting the location of the stabilized k-fiber region one timepoint prior to fracture. **h)** The distribution of k-fiber stabilization outcomes based on position of manipulation site at fracture relative to initial manipulation site.

To ask whether local force was sufficient to stabilize the lattice prior to fracture, we pulled a k-fiber for ∼2 minutes at 4 µm/min, removed the microneedle and rapidly laser ablated that same k-fiber between the manipulation site and the kinetochore (Figure 2c-d, Video 4). While this is a technically challenging experiment for several reasons, in 100% of the six cases where we successfully ablated the k-fiber after manipulation, the k-fiber resisted depolymerization at or near the manipulation site for tens of seconds to minutes (Figure 2e-f). Yet, microneedle pulling alone only led to stabilization of about 50% of new k-fiber plus-end stubs (Figure 1f, 2e). We hypothesized that the lattice is only stabilized locally and that the position of the fracture site with respect to the force application site determines the odds of stabilization. Consistent with this, stabilization was much more likely if the needle slid away from, rather than toward, the pole during manipulation (Figure 2g-h). Thus, if the fracture site (microneedle or laser ablation induced) is towards the plus-end relative to the force application site, stabilization is more likely. Importantly, in all cases where the microneedle was present along the k-fiber for ≥ 2 minutes without moving we saw rapid k-fiber depolymerization upon ablation (Figure 2e-f). Thus, the microneedle alone is not sufficient to stabilize the k-fiber. Together, we conclude that applied load is both necessary and sufficient for spindle microtubule lattice stabilization, independent of microtubule fracture.

### New spindle microtubule plus-ends created by local load have arrested dynamics

To gain insight into how force may have changed k-fiber microtubules at the fracture site, rather than at kinetochore-bound plus-ends, we sought to measure the dynamics of new plus-ends after load-induced fracture. To do so, we created a bleach mark on the fractured k-fiber stub pole-side (Figure 3a-b, Video 5) and tracked its position along the stub over time. The distance between the mark and newly formed plus-ends remained nearly stable: it depolymerized at only 0.28 ± 0.15 µm/min (Figure 3c-e, Supplementary Figure 2), two orders of magnitude slower than depolymerization triggered by acutely laser severing k-fibers (∼20 µm/min)^41^. We also never observed spindle microtubule growth from the fractured plus-ends. However, the average mitotic microtubule polymerization speed is ∼13 µm/min^44^, again two orders of magnitude greater than the measured plus-end depolymerization rate we measure. Together, this data shows that sustained local load can fracture spindle microtubules and create new plus-ends with arrested dynamics, neither polymerizing nor depolymerizing at expected rates.

**Figure 3:**
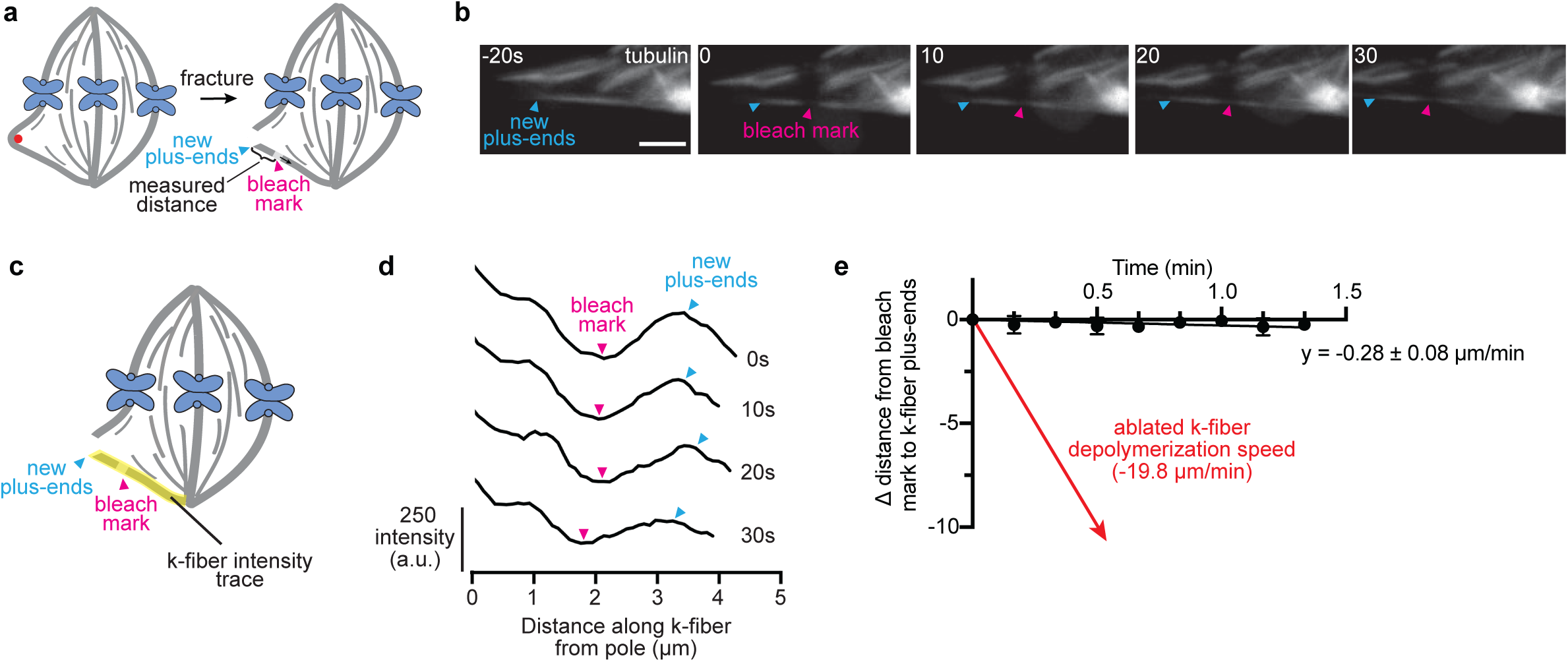
New spindle microtubule plus-ends created by local load have arrested dynamics. **a)** Schematic depicting microneedle load-induced fracture, and k-fiber stub bleaching (pink arrowhead) post-fracture to probe the dynamics of new plus-ends (blue arrowhead) after local load application. We measure the distance between the plus-ends (blue arrowhead) and bleach mark (pink arrowhead). **b)** Representative timelapse of a fractured k-fiber stub showing the generation of a bleach mark (at time 0:00) immediately post-fracture and subsequent tracking of the new plus-ends (blue arrowhead) and bleach mark (pink arrowhead) over time in PtK2 cells stably overexpressing GFP-tubulin (white). Scale bar = 2 µm. **c)** Schematic depicting a fractured k-fiber with a tubulin intensity trace along the k-fiber stub attached to the pole. The tubulin intensities were used to determine the location of the bleach mark and new plus-ends relative to the spindle pole. **d)** Representative traces of the fractured k-fiber pole-side stub tubulin intensity along its length. The locations of the bleach mark (pink arrowhead) and new plus-ends (blue arrowhead) are denoted. **e)** The change in distance between the newly formed plus-ends and the bleach mark plotted over time compared against known k-fiber depolymerization speed upon laser-induced severing^41^ (n=5 k-fibers, N=2 experimental days slope calculated using linear regression analysis, slope tested for significant difference from 0, p<0.01).

### Local force-induced spindle microtubule curvature results in a local reduction of doublecortin

One mechanism by which locally applied force could change the microtubule lattice is by changing the spacing between tubulin dimers in the lattice^45^. To test this possibility and define how far from the load application site such lattice spacing change could occur, we used doublecortin (DCX) as a proxy for microtubule lattice spacing. Doublecortin is thought to preferentially bind compressed portions of the microtubule lattice containing GDP-tubulin dimers^46^. We again employed microneedle manipulation to apply local loads, this time in PtK2 cells stably overexpressing HaloTag-tubulin and transiently overexpressing eGFP-DCX (Figure 4a-b, Video 6). We quantified the DCX:tubulin intensity ratio and radius of curvature of the manipulated k-fiber throughout the manipulation.

**Figure 4:**
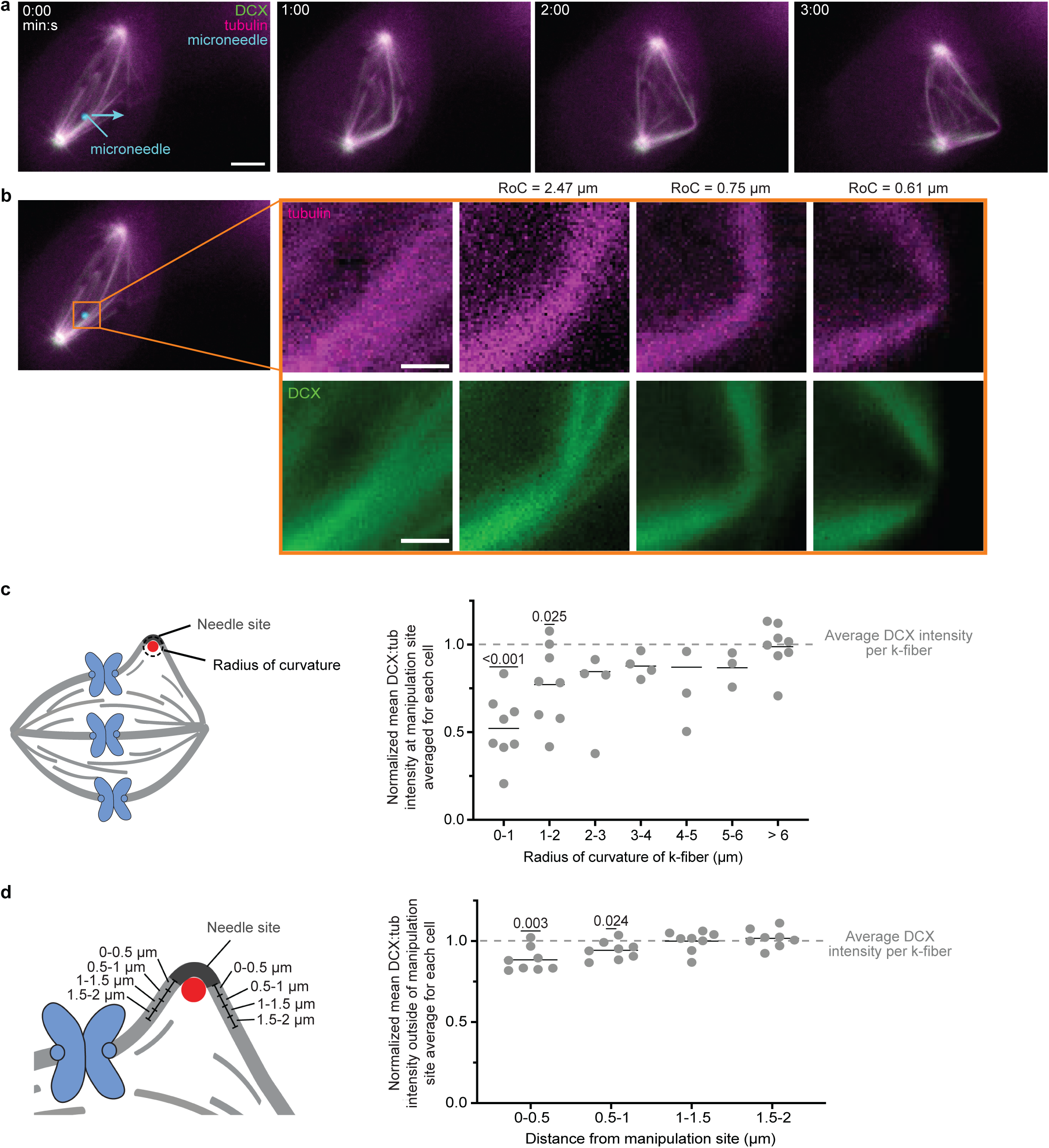
Local force-induced spindle microtubule curvature results in a local reduction of doublecortin. **a)** Representative timelapse of PtK2 cells stably overexpressing HaloTag-tubulin (magenta, JF 646) and transiently overexpressing eGFP-doublecortin (green) where a metaphase k-fiber is subject to microneedle (cyan, AF561 labelled) manipulation over 15 μm (blue arrow direction) in 216 s starting at time 0:00 (4 μm/min manipulation speed). Scale bar = 5 µm. **b)** Zoomed in timelapse of the manipulated region of the k-fiber with the measured k-fiber radius of curvature (RoC) displayed and the tubulin and doublecortin channels separated. Scale bars = 1 µm. **c)** The mean doublecortin:tubulin intensity ratio along the curved portion wrapped around the microneedle normalized to the mean doublecortin:tubulin intensity ratio of the whole manipulated k-fiber binned by radius of curvature (n=20 k-fibers, N=4 experimental days, one sample t and Wilcoxon test, mean tested for significant difference from 1). **d)** The mean doublecortin:tubulin intensity ratio of 0.5 μm sections of manipulated k-fibers for all timepoints where the k-fiber’s radius of curvature is <1 μm, binned by distance from the manipulation site (n=32 measurements from 8 k-fibers, each point represents the average intensity ratio within the specified region for a given k-fiber, one sample t and Wilcoxon test, mean tested for significant difference from 1).

We found that the DCX:tubulin intensity ratio is lower at smaller radii of curvature at the manipulation site (Figure 4c), consistent with observations in single microtubules with naturally occurring curvature at interphase^46^. Specifically, we observed a significant reduction in the DCX:tubulin intensity ratio at the manipulation site when k-fibers reached radii of curvature <2µm (Figure 4c). To determine if this lattice structural change propagated along manipulated k-fibers, we examined cases where a radius of curvature of <1µm was achieved. In these cases, a slight, but statistically significant reduction in DCX:tubulin intensity ratio was found up to 1 µm from the manipulation site (Figure 4d). Thus, sustained local load causes a loss of doublecortin at and near sites of high microtubule curvature. Together, this indicates that the local loads we apply in the spindle locally remodel the k-fiber microtubule lattice in a manner consistent with lattice stretching and/or GTP-tubulin integration.

### Local force leads to local EB1 recruitment along spindle microtubule lattice before fracture and at new plus-ends after fracture

Together, reduced DCX (Figure 4) and increased lattice stability (Figures 1-3) under local load are consistent with GTP-tubulin integration at and near the force application site. In the absence of a live-cell imaging method to directly discriminate between GTP-and GDP-tubulin, we used end-binding protein 1 (EB1) as a proxy for GTP-tubulin. EB1 is believed to preferentially bind to GTP-tubulin in a nucleotide-dependent manner^47,48^. We used PtK1 cells stably expressing GFP-EB1 and visualized microtubules with a low concentration of SiR tubulin, which is not expected to influence spindle microtubule dynamics^49^. If GTP-tubulin integration is responsible for increased spindle microtubule stability under load, we expect EB1 to be locally enriched at sites along the microtubule lattice.

To test this prediction, we performed microneedle manipulation on k-fibers and examined the EB1:tubulin intensity ratio at stable plus-ends upon k-fiber fracture (Figure 5a-b, Video 7). We compared this ratio to two other regions of the manipulated k-fiber: near the kinetochore and over the rest of the k-fiber, which we termed ‘bulk k-fiber’ (Figure 5c). We found that EB1:tubulin intensity is significantly enriched at the newly formed plus-ends upon fracture (Figure 5d). We also compared the EB1:tubulin intensity ratio of the manipulated k-fiber to a control k-fiber in the same cell that was not manipulated to determine if EB1 was enriched locally or globally along the manipulated k-fiber (Figure 5e). We found that there was no significant difference in EB1 enrichment along the’bulk’ portion of the manipulated k-fiber compared to the control, further confirming that the effect of load is local. We repeated these experiments with PtK2 cells stably overexpressing HaloTag-tubulin and transiently overexpressing 2x-GFP-EB1 to exclude the possibility of SiR tubulin affecting EB1 localization and made consistent observations (Supplemental Figure 3). These data support the hypothesis that GTP-tubulin is present at newly formed plus-ends post fracture and gives rise to their stabilization.

**Figure 5:**
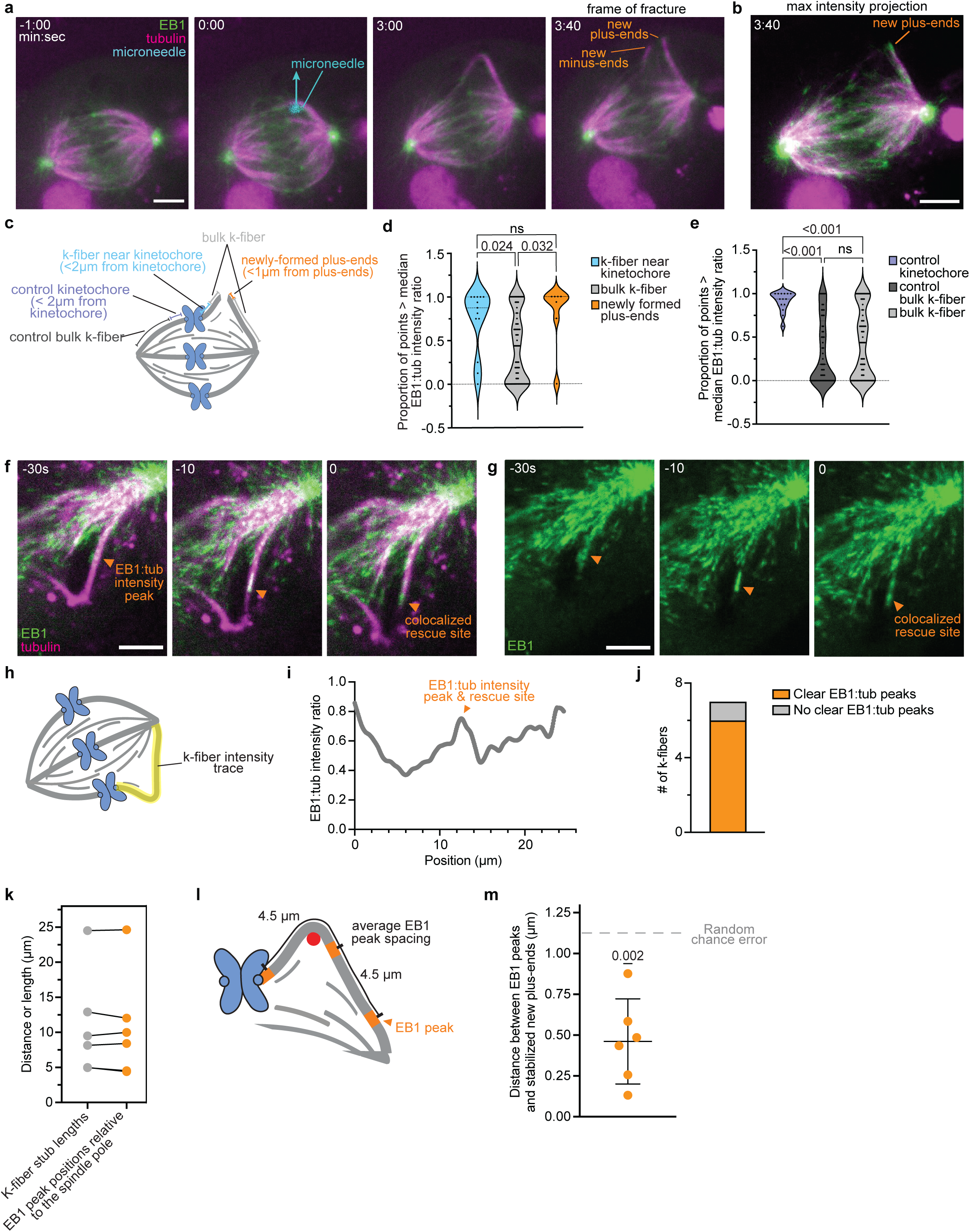
Local force leads to local EB1 recruitment along spindle microtubule lattice before fracture and at new plus-ends after fracture. **a)** Representative timelapse of a PtK1 cell stably overexpressing GFP-EB1 (green), treated with SiR tubulin (magenta), where a metaphase k-fiber is subject to microneedle (cyan, AF561 labelled) manipulation of 30 μm over 216 s (8 μm/min pull speed) until fracture. New minus-ends and plus-ends are marked (orange). Scale bar = 5 µm. **b)** Maximum intensity projection of the representative cell in (a) ∼30 s post-fracture. Scale bar = 5 µm. **c)** Schematic depicting different regions within the manipulated spindle where the EB1:tubulin intensity ratio was quantified. **d & e)** Distribution of the percentage of points greater than the median EB1:tubulin intensity ratio along the manipulated and control k-fibers binned by location d) (N=7 n_k-fiber near kinetochore_=13, n_bulk k-fiber_=85, n_newly formed plus-ends_=7 1-µm sections, M=3 experimental days, ordinary one-way ANOVA). e) (N=7 k-fibers, n_control kinetochore_=14, n_control bulk k-fiber_=61, n_bulk manipulated k-fiber_= 85 1-µm sections, M=3 experimental days, ordinary one-way ANOVA). **f & g)** Representative timelapse of a PtK1 cell stably overexpressing GFP-EB1 (green), treated with SiR tubulin (magenta), where a metaphase k-fiber is subject to microneedle manipulation of 30 μm over 216 s (8 μm/min pull speed) until fracture. Site of EB1 enrichment is marked (orange). Panel g only contains the GFP-EB1 channel (green). Scale bars = 5 µm. **h)** Schematic depicting trace used to determine the EB1:tubulin intensity ratio along the manipulated k-fiber prior to fracture. **i)** Representative EB1:tubulin intensity ratio along the length of the k-fiber shown in panels f & g. **j)** Distribution of the number of cells that demonstrated ≥1 EB1:tubulin intensity peak along the length of the manipulated k-fiber one timepoint prior to fracture. **k)** Plot of the measured length of the k-fiber post fracture compared to the closest EB1:tubulin peak position one frame prior to fracture (n=6 k-fibers, N=3 experimental days). **l)** Schematic depicting the mean EB1 peak frequency (assuming peaks are equally spaced) along a k-fiber, based on the measured EB1:tubulin intensity data from k-fibers demonstrating ≥1 peak. **m)** The distance between the length of the k-fiber post-fracture to the closest EB1:tubulin peak position one frame prior to fracture (N=6, M=3 experimental days, one sample t-test, tested for significant difference from the expected value of 1.125 µm)

Finally, we sought to determine whether EB1 only bound to new plus-ends after they were created, or also to the microtubule lattice pre-facture. The latter is expected if GTP-tubulin integration led to lattice stabilization pre-fracture (Figure 2). Upon inspection (Figure 5f-g, Video 8), we found that many (6/7) of the manipulated k-fibers exhibited distinct EB1 intensity peaks along their length prior to fracture (Figure 5h-j). Further, we found that the length of the stabilized k-fiber stub corresponded well with an EB1 peak location in all cases (Figure 5k). In contrast, if EB1 peaks were randomly distributed we expect them to occur every 4.5 µm (based on the total length of manipulated k-fibers divided by the number of EB1 peaks detected before fracture) (Figure 5l). Given this, the closest a point could be to an EB1 peak is 0 µm, and the furthest is 2.25 µm. Thus, if stabilization sites did not correlate with the location of EB1 peaks we expect the mean difference between peak and stabilization locations to be ∼1.125 µm (the mean of the closest and furthest distances). Instead, we found that the mean difference was significantly less (0.46 ± 0.26 µm) (Figure 5m). Together, these findings are consistent with local load causing local spindle microtubule lattice damage that is repaired prior to fracture through the integration of GTP-tubulin, giving rise to spindle microtubule stabilization (Figure 6).

**Figure 6:**
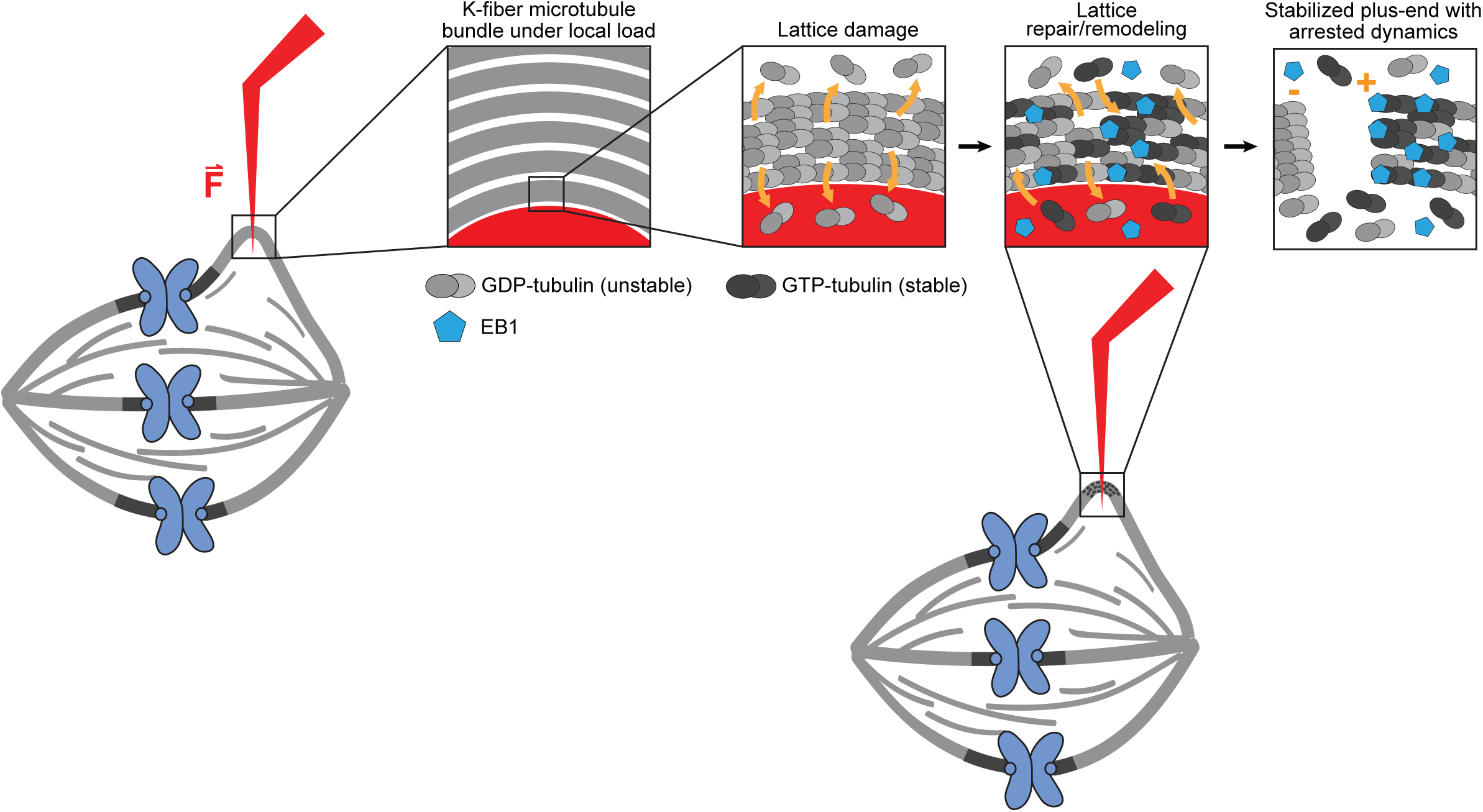
Model for spindle microtubule damage, remodeling and stabilization under sustained local load. Left: Sustained local load from microneedle (red) pulling on a k-fiber microtubule bundle leads to gradual loss (orange dashed arrows) of individual GDP-tubulin dimers (light grey dimers) from individual protofilaments of individual microtubules (left). Local microtubule lattice stretching under load (Figure 4) makes GDP-tubulin dimer loss more likely by weakening dimer-dimer contacts. These lattice vacancies are then repaired via integration (orange arrows) of freely diffusing GTP-tubulin dimers (dark grey dimers), causing EB1 (blue pentagon) lattice localization (Figure 5) and lattice stabilization (Figures 1-3). We propose that force-induced k-fiber microtubule lattice stabilization helps reinforce the spindle where it is under critical loads. Right: The new plus-end is composed of a mixed nucleotide zone due to partial damage and thus partial remodeling, resulting in a stabilized plus-end with arrested dynamics upon fracture (Figure 3).

## Discussion

The ability of the spindle to maintain its structure in the face of mitotic and environment forces is crucial to proper chromosome segregation. Here, we use microneedle manipulation to mechanically challenge the mammalian spindle and examine how the lattice of spindle microtubules responds to local load. We show that sustained local load results in k-fiber fracture and microtubule stabilization, with new plus-ends having arrested dynamics (Figures 1-3). We demonstrate that this load locally changes spindle microtubule lattice structure and biochemistry (Figures 4-5), via remodeling consistent with local GTP-tubulin incorporation along the lattice, near the force application site (Figure 5). Together, we propose that force-induced microtubule lattice damage, remodeling and stabilization helps locally reinforce the spindle when and where it is under critical load (Figure 6).

We have long known that mechanical force increases the stability of kinetochore-microtubule attachments^22,50^, and regulates the end dynamics of kinetochore-microtubules^16,26,51^. Here, we show that mechanical force not only regulates the ends of spindle microtubules but can locally regulate microtubule stability and lattice remodeling – where load is applied (Figures 1, 3, 4, and 5). We show that spindle microtubule stability can also be governed by a microtubule intrinsic mechanism acting along the length of microtubules, not just at their ends (Figures 1, 2, and 5).We propose that the applied load creates tension on the microtubule lattice, weakening interactions between tubulin dimers and thus increasing the likelihood that dimers dissociate from the lattice, leaving lattice vacancies that can be repaired (Figure 6). Our ability to exert local loads along spindle microtubules revealed highly local structural and biochemical responses to force (Figures 1, 2, 4, and 5), which we propose underlies the variability in newly formed plus-end stabilization seen in load-induced fracture events (Figure 1f): in this model, the location of the fracture site relative to the repair site dictates whether stabilization occurs. In principle, spindle microtubules responding to force all along their length, and not just at ends, could provide greater tunability over the spatial distribution of spindle microtubule stability and dynamics, and allow passive reinforcement of spindle regions under critical loads.

Upon fracture of k-fiber microtubules (Figure 1), new microtubule plus-ends exhibit arrested dynamics (Figure 3), depolymerizing almost two orders of magnitude slower than k-fiber plus-ends acutely created by laser ablation^41^. How to think of these arrested dynamics given the long-held dynamic instability model^52^, where plus-ends rapidly either grow or shrink, is not yet clear. The observed plus-end depolymerization speed difference matches differences between GTP-and GDP-tubulin dissociation rates^53,54^, suggesting new plus-ends created by mechanical force are biochemically distinct from those created by laser ablation. It remains unclear how fractured k-fiber microtubules maintain arrested dynamics for tens of seconds to minutes, and if this observation relates to prolonged slow growth at interphase^55^ or a metastable plus-end state seen *in vitro*^56^. We favor a model whereby the microtubule plus-end is composed of a mixed nucleotide zone^57^, where GTP-and GDP-tubulin dimers are both present, resulting in de/polymerization incompetent microtubules (Figure 6). Alternatively, a collection of microtubule-associated proteins (MAPs) at plus-ends, such as CLASPs^58,59^, kinesin-4^60^, SSNA1^61^, or CSPP1^62^, among others, could give rise to arrested dynamics, as seen in suppression of plus-end dynamics *in vitro*. The development of tools to distinguish between tubulin nucleotide states in live cells will be key to determining the mechanism behind force-induced spindle microtubule stabilization and arrested dynamics.

Microtubule lattice repair was first reported decades ago^63^. It has since been studied *in vitro*^32,35,36^ and more recently *in vivo* at interphase^33,64^. We now know that microtubules subject to bending stress or motor traffic accrue damage along their length that can be repaired^32,35,64–66^, and that lattice repair sites, termed “GTP islands”, serve as rescue sites when microtubules depolymerize^33,67^. Unlike *in vitro* or interphase microtubules, however, k-fiber microtubules exist in a bundle. Thus, many microtubules must be repaired at largely the same time and place (see splayed ends, Figure 1i) to give rise to whole k-fiber stabilization. Whether kinetochore-microtubules are independently damaged and remodeled, or whether these occur at the level of a whole bundle, is an open question for future study. In principle, after a first spindle microtubule in a bundle breaks under force, remaining microtubules are increasingly likely to break as fewer microtubules now bear the same load, providing cooperativity to the k-fiber fracture process.

Looking forward, our work opens the question of how force leads to spindle microtubule lattice damage, whether and how MAPs^37,61,62^ aid in damage repair and stabilization, and what function this remodeling provides. For example, akin to interphase ^68^ global force from the environment could lead to global lattice remodeling and spindle reinforcement. Alternatively, lattice remodeling could help repair more local spindle damage: k-fiber microtubules are often curved, locally under tension and trafficked by motor proteins, all of which could increase, perhaps synergistically, the likelihood of lattice damage and need for repair and stabilization to maintain spindle structure. More broadly, our work motivates us to probe this phenomenon across evolution and other microtubule structures, particularly those with physically challenging environments or functions.

## Acknowledgements

We thank members of the Dumont Lab for helpful discussions and critical reading of the manuscript. We thank Torsten Wittman for helpful discussions and the EB1 and doublecortin (DCX) constructs. We thank Alexey Khodjakov for the PtK2 GFP–α-tubulin cell line and Ted Salmon for the PtK1 GFP-EB1 cell line. We thank Pooja Suresh and Alex Long for technical training and helpful conversations. This work was supported by the NSF Graduate Research Fellowship Program (C.J.R.), UCSF Discovery Fellowship (C.J.R), UC Berkeley/UCSF Bioengineering Community Fellowship (C.J.R), H2H8 Research Grant (C.J.R.), NIH R35GM136420 (S.D.) and NSF 1548297 (S.D). S.D. is a Chan Zuckerberg Biohub investigator.

## Author Contributions

Conceptualization, C.J.R., M.K.C., S.D.; methodology, C.J.R.; software, C.J.R.; validation, C.J.R.; formal analysis, C.J.R.; investigation, C.J.R., V.M.; resources, C.J.R., M.K.C., N.H.C., S.D.; data curation, C.J.R.; writing – original draft, C.J.R.; writing – review and editing, C.J.R., M.K.C., V.M., N.H.C., and S.D.; visualization, C.J.R.; supervision, S.D.; funding acquisition, C.J.R. and S.D.

## Methods

### Cell Culture

PtK2 cells were cultured in MEM (Invitrogen) supplemented with sodium pyruvate (Invitrogen), nonessential amino acids (Invitrogen), penicillin/streptomycin, and 10% qualified and heat-inactivated fetal bovine serum (Invitrogen) and maintained at 37°C and 5% CO_2_. PtK2 cells stably expressing human GFP–α-tubulin (gift from Alexey Khodjakov, Wadsworth Center, Albany, NY) and PtK2 cells stably expressing human β-tubulin-HaloTag were used^69^. PtK1 cells were cultured in Ham’s F12 media (Invitrogen) supplemented with penicillin/streptomycin and 10% qualified and heat-inactivated fetal bovine serum and maintained at 37°C and 5% CO2. PtK1 cells stably expressing human GFP-EB1 were used (gift from Ted Salmon, UNC Chapel Hill, Chapel Hill, NC).

### Transfection Plasmids

PtK2 cells stably expressing human β-tubulin-HaloTag were transfected using Viafect (Promega) and imaged 48h after transfection with 2x-GFP-EB1 or eGFP-DCX plasmids (both gifts from Torsten Wittman, UCSF, San Francisco, CA).

### Dyes

PtK2 cells stably expressing β-tubulin-HaloTag were treated with 100 nM Janelia Fluor 646 (GA1120, Promega) for 1 h before imaging. PtK1 GFP-EB1 cells were incubated with SiR-tubulin at 100 nM and verapamil at 10 µM (Cytoskeleton, Inc.) for 1 h before imaging. This concentration of SiR-tubulin was previously shown not to impact spindle microtubule dynamics^30^.

### Microscopy

For live cell imaging, cells were plated onto #1.5 glass-bottom 35 mm dishes coated with poly-D-lysine (P35G-1.5-20-C, MatTek Life Sciences). Live cells were imaged using an inverted microscope (Eclipse Ti-E; Nikon) with a spinning disk confocal (CSU-X1; Yokogawa), head dichroic Semrock Di01-T405/488/568/647 for multicolor imaging, equipped with 405 nm (100 mW), 488 nm (120 mW), 561 nm (150 mW), and 642 nm (100 mW) diode lasers, emission filters ET455/50M, ET525/50M, ET630/75M, and ET690/50M for multicolor imaging, and either an iXon3 camera (Andor Technology) or a Zyla 4.2 sCMOS camera (Andor Technology) operated by MicroManager 2.0.3. Images were acquired with a 100x 1.45 Ph3 oil objective (Nikon Instruments) with the exception of Figure 3 data, which was acquired with a 60x 1.40 Ph3 oil objective. Cells were imaged in a stage-top incubation chamber (Tokai Hit) with the top lid removed and maintained at 30°C.

### Microneedle Manipulation and Microneedle Preparation

Microneedle manipulation and needle fabrication was performed as outlined in previous work^15,30^. Microneedles were made from glass capillaries with an inner and outer diameter of 1 mm and 0.58 mm respectively (1B100-4 or 1B100F-4, World Precision Instruments). Glass capillaries were pulled with a micropipette puller (P-87, Sutter Instruments), bent and polished using a microforge (Narishige International) according to the same specifications, parameters, and geometries described earlier^15^. These parameters allowed for the needle to approach cells orthogonal to the imaging plane and to conduct manipulations without rupturing the cell^15^. Prior to imaging, microneedles were coated with BSA-Alexa-555 (A34786, Invitrogen) or BSA-Alexa-647 (A-34785, Invitrogen) by soaking them in coating solution for 60 s. Coating solution was obtained by dissolving BSA-Alexa dye and Sodium Azide (Nacalai Tesque) in 0.1 M phosphate-buffered saline (PBS) at a final concentration of 0.02% and 3 mM, respectively^70^. This coating solution allowed needles to be visualized via fluorescence imaging, aiding in positioning of the needle along a single k-fiber.

The micromanipulator was mounted to the microscope body and positioned above samples^15^. Manipulations were performed in 3D using a x-y-z stepper-motor micromanipulator (MP-225, Sutter Instruments). A 3-axis-knob (ROE-200, Sutter Instruments) was connected to the manipulator via a controller box (MPC-200, Sutter Instruments). In all manipulations, prior to manipulating the needle was positioned via phase imaging at the approximate x-y position of the intended manipulation site ∼ 10 µm above the cell in z. While imaging every 10 s at a single z-plane, the needle was manually lowered into place along a single k-fiber. If necessary, the needle’s position in x-y was changed by raising the needle above the cell and repositioned in x-y before approaching the intended k-fiber. Once properly positioned, a Python script^15^ was used to communicate with the Sutter Multi-link software (Multi-Link, Sutter Instruments) to move the microneedle a defined distance (62.5 nm resolution) approximately orthogonal to the pole-pole axis over a defined period of time (0.9 s time resolution). This computer-controlled movement of the microneedle allowed for consistent, reproducible experiments across all studied conditions.

We analyzed manipulated cells demonstrating all listed attributes: cell health was not significantly impacted by manipulation (no cell rupture or significant damage to the entirety of the spindle), the manipulated k-fiber was a part of a mature metaphase spindle (with chromosomes forming a tight metaphase plate), and k-fibers could be tracked during and after manipulation to determine fracture (or no fracture), stability (or lack of stability) and/or protein localization along the manipulated k-fiber.

### K-fiber Fracture Experiments (Figure 1)

Individual metaphase k-fibers were targeted for microneedle manipulation over distances of 15, 21, and 30 µm over 216 s, resulting in approximate manipulation speeds of 4, 6, and 8 µm/min, respectively. Manipulations were performed while imaging every 10 s at a single z-plane. Upon completion of needle movement, the needle remained in place until fracture occurred or >7 min passed since the start of the manipulation. Upon fracture, the current acquisition was stopped and the needle removed before capturing a full cell volume (± 5 µm, 0.3 µm spacing) to verify the k-fiber indeed fractured – as determined by no microtubule signal beyond background connecting the k-fiber stubs and a clear mechanical uncoupling between k-fiber stubs (i.e. both stubs taking dramatically different orientations after fracture events).

### Laser Ablation (Figure 2)

Laser ablation experiments were done using 514 nm ns-pulse laser light and a galvo-controlled MicroPoint Laser System (Andor, Oxford Instruments) controlled through MicroManager 2.0.3. Prior to each experiment, the ablation laser was aligned in the x-, y-and z-axes. While imaging every 10 s at a single z-plane, k-fibers were subject to microneedle manipulation at the 4 µm/min manipulation speed for ∼ 2 min. Manipulated k-fiber ablations were performed immediately after microneedle removal by firing the laser at 2–3 discrete points (60–90 pulses of 3 ns at 20 Hz), along a small area of the manipulated k-fiber between the kinetochore and manipulation site. Mechanical uncoupling (i.e. both stubs taking dramatically different orientations after fracture events). of the two k-fiber stubs was used to indicate a successful ablation. The same parameters were used for control, unperturbed k-fibers except imaging was performed at higher time resolution (2 s).

### Photobleaching (Figure 3)

Photobleaching experiments were done using a UGA-42 Firefly (Rapp OptoElectronic, Hamburg/Germany) photomanipulation system outfitted with a 473 nm, 100 mW laser. The UGA-42 Firefly was controlled via SysCon 2 (2.6.0) software. Individual metaphase k-fibers were subject to microneedle manipulation at 4 µm/min pull speeds while imaging a single z-plane every 10 s. Upon fracture, the needle was removed, and multiple z-planes (5) were imaged (± 1 µm, 0.5 µm spacing) every 10s. An area along the length of the k-fiber between the newly formed plus-end and spindle pole was stimulated via the UGA-42 Firefly to induce a bleach mark to track k-fiber microtubule dynamics.

### DCX Localization (Figure 4)

Individual metaphase k-fibers in HaloTag-tubulin PtK2 cells expressing eGFP-DCX were targeted for microneedle manipulation. K-fibers were subject to manipulation at 8 µm/min manipulation speed. Manipulations were performed while imaging every 10 s at a single z-plane.

### EB1 Localization (Figure 5)

Individual metaphase k-fibers GFP-EB1 PtK1 cells expressing were targeted for microneedle manipulation. K-fibers were subject to manipulation at 8 µm/min manipulation speed. Manipulations were performed while imaging every 10 s at a single z-plane. Upon fracture, the current acquisition was stopped and the needle removed before capturing a full cell volume (± 5 µm, 0.3 µm spacing). The same protocol was repeated HaloTag-tubulin PtK2 cells transiently expressing 2x-GFP-EB1, however the EB1 channel was imaged only after fracture.

### Image Analysis

All image analysis was performed in Fiji (2.14.0). In some cases, data obtained from Fiji was transferred to MATLAB (R2022a) for further processing and quantification.

#### K-fiber fracture and time to fracture measurements (Figures 1e,j,k)

K-fiber fracture was determined through a combination of multiple factors including:

- Loss of tubulin signal (above background) between the k-fiber stubs, initially seen in single-z plane imaging, confirmed through volumetric imaging upon suspected fracture;
- Mechanical uncoupling of the k-fiber stubs, resulting in stubs taking on dramatically different orientations;
- Rapid change of plane of one or both k-fiber stubs, indicating tension across the k-fiber was likely lost due to fracture.

Time to fracture was calculated based upon the time between the first frame with microneedle computer-controlled movement and when the requirements for fracture were met (listed above).

#### K-fiber stability (Figures 1f and 2f)

The full cell volume captured ∼30 s after suspected k-fiber fracture was used to determine k-fiber plus-end stability. If a k-fiber stub attached to the spindle pole was present (indicating it did not rapidly depolymerize), the plus-end was determined to be stabilized. In the case of laser ablation with or without microneedle manipulation, stability was determined through single-z plane imaging. If a k-fiber stub attached to the spindle pole remained for >30 s, it was deemed stable.

#### K-fiber length change measurements (Figure 1g, Supplementary Figure 1)

The initial k-fiber length and control k-fiber length were measured at the start of manipulation. One timepoint prior to fracture, the manipulated and control k-fiber lengths were manually measured. In Figure 1g, the calculated k-fiber length change was compared to the distance of the fracture site along the k-fiber relative to the kinetochore. Fracture sites located a greater distance from the kinetochore than the measured k-fiber length change were deemed to be stable sites located outside of the newly polymerized tubulin region.

#### K-fiber intensity measurements (Figure 1h)

Two traces orthogonal to the manipulated k-fiber (one between the microneedle and the kinetochore and another between the microneedle and the spindle pole) were obtained to determine the overall k-fiber intensity at the beginning of microneedle manipulation and one timepoint prior to k-fiber fracture. Additionally, a trace was generated orthogonal to an unmanipulated control k-fiber in the same cell at the same two timepoints to control for bleaching. Once obtained, a custom MATLAB code was used to calculate the area underneath the FWHM region of the intensity curves to yield the integrated intensity. An additional operation was performed to remove any area under the curve that was a result of background noise (causing the peak to be offset from 0). The values were normalized according to the final integrated intensity/initial integrated intensity for each of the three regions measured.

#### K-fiber radius of curvature (Figures 1l and 4c)

The k-fiber radius of curvature was measured by manually fitting a circular trace to the k-fiber signal at the site of manipulation.

#### Microneedle position along k-fiber (Figure 2g)

The distance between the microneedle and spindle pole (along the k-fiber) was measured at the start of manipulation and one frame prior to fracture. If the measured distance increased, it was classified as “further from the pole”; if it decreased, it was classified as “closer to the pole”; if it remained the same, it was classified as “same distance from pole.”

#### Bleach mark position and k-fiber length (Figure 3d-e, Supplementary Figure 2)

An intensity trace of the fractured k-fiber stub from the pole to beyond the newly formed plus-end was manually generated for each timepoint. A custom MATLAB script was used to identify the location of the bleach mark and the end of the k-fiber stub. Intensity data was inverted, and a peak detection algorithm was used to find the center of the bleach mark along the k-fiber stub length at each timepoint. To determine the end of the k-fiber stub, the intensity data along the last 20 points prior the end were examined and the location of the greatest change in intensity (high to low) was used to define the k-fiber stub end as determined through the MATLAB ‘findchangepts’ function.

#### DCX:tubulin intensity ratio (Figure 4c-d)

An intensity trace of the manipulated k-fiber was performed at each timepoint during the manipulation. The tubulin, DCX, and needle intensities were obtained. Using a custom MATLAB code, the location and width (based on FWHM) of the microneedle was determined tusing the microneedle intensity data and a peak detection algorithm. This region of the k-fiber was deemed the ‘manipulation site.’ The mean DCX:tub intensity ratio for this region was normalized to the mean DCX:tub intensity of the k-fiber outside the manipulation site (Figure 4c). Additionally, for timepoints where the radius of curvature of the manipulation site was < 1 µm, the k-fiber was discretized into a series 0.5 µm sections on each side of the manipulation site and their relative location compared to the manipulation site saved. The mean DCX:tub intensity for 0.5 µm sections were normalized to the mean DCX:tub intensity of the k-fiber outside the manipulation site (Figure 4d). In both cases, the data was aggregated for each individual k-fiber at various timepoints and the mean displayed and used for statistical tests.

#### EB1:tubulin intensity ratio (Figure 5d-e, Supplementary Figure 3b)

An intensity trace of the manipulated k-fiber and a control k-fiber in the same cell was performed at the first timepoint after fracture after background correction was performed (average background of three 10×10 pixels boxes within the manipulated cell away from the spindle structure). A custom MATLAB code was used to determine the end of each side of the k-fiber stub for the manipulated k-fiber and the start of the k-fiber relative to the kinetochore (all based on tubulin signal). Based on these locations, the fractured k-fiber data was binned into three regions (newly formed plus-end {< 1 µm from the k-fiber stub end}, k-fiber near kinetochore {< 2 µm from the k-fiber original plus-end}, and the rest, bulk k-fiber) and the mean EB1:tub intensity ratio of those regions calculated and used for statistical tests. The control k-fiber was also binned into two regions (control kinetochore {< 2 µm from the k-fiber plus-end} and the rest, bulk control) and the mean EB1:tub intensity ratio of those regions calculated and used for statistical tests (Figure 5d, 5e, and Supplementary Figure 3).

#### EB1 peak sites and rescue locations (Figure 5k and m)

An intensity trace of the manipulated k-fiber in the last timepoint where the entire k-fiber was in plane prior to fracture was obtained. Using a custom MATLAB code, the EB1:tub intensity ratio along the entire k-fiber length was obtained. After smoothing the data, MATLAB’s ‘findpeaks’ function was used to determine number of EB1:tub peaks and their respective location. The length of the fractured k-fiber stub attached to the pole at the timepoint of fracture was determined based on tubulin signal. This length was compared to the closest EB1:tub peak for further statistical analysis.

## Statistical analysis

Details of statistical tests and sample sizes (number of cells and number of independent experiments, defined by individual imaging sessions performed on different days) are provided in figure legends. Two-sided paired t-tests were used to compare continuous datasets within the same experiment. Ordinary one-way ANOVAs were used to compare continuous datasets across experiments containing more than two categories. Simple linear regressions were fitted to continuous data sets containing one independent and one dependent variable. One sample t-tests were performed on continuous datasets that were normalized or compared against a known measure indicated in figure legends. We used p < 0.05 as the threshold for statistical significance.

## Video Legends

**Video 1 – Figure 1 panel c - Microneedle manipulation of individual metaphase k-fiber until fracture with lack of plus-end stabilization:** Microneedle manipulation of individual k-fiber reveals an example of lack of newly-formed plus-end stabilization upon fracture in representative PtK2 cell stably overexpressing GFP-tubulin. The microneedle (red, AF647 labelled) exerts force on the k-fiber (white) and is subsequently removed upon fracture and a full cell volume is acquired (0 ± 4 µm, 0.3 µm z-spacing) to create a maximum intensity projection. Scale bar, 5 µm. Video was collected using a spinning disk confocal microscope at one frame every 10 s. Video was stopped once fracture occurred. Video has been adjusted to play back at a constant rate of 5 frames/s. Video corresponds to still images from Fig. 1 panel c.

**Video 2 – Figure 1 panel d - Microneedle manipulation of individual metaphase k-fiber until fracture and subsequent plus-end stabilization:** Microneedle manipulation of individual k-fiber reveals an example of newly-formed plus-ends stabilized upon fracture in representative PtK2 cell stably overexpressing GFP-tubulin. The microneedle (red, AF647 labelled) exerts force on the k-fiber (white) and is subsequently removed upon fracture and a full cell volume is acquired (0 ± 4 µm, 0.3 µm z-spacing) to create a maximum intensity projection. Video was stopped once fracture occurred. Scale bar, 5 µm. Video was collected using a spinning disk confocal microscope at one frame every 10 s. Video has been adjusted to play back at a constant rate of 5 frames/s. Video corresponds to still images from Fig. 1 panel d.

**Video 3 – Figure 2 panel a – Laser ablation of an unperturbed metaphase k-fiber:** Laser ablation of an unperturbed individual k-fiber reveals the lack of stability of the newly formed plus-end in representative PtK2 cell stably overexpressing GFP-tubulin. The k-fiber (white) is ablated (yellow “X”) and its newly formed plus-end tracked (blue arrow), showing rapid depolymerization to the spindle pole. Scale bar, 5 µm. Video was collected using a spinning disk confocal microscope at one frame every 10 s. Video has been adjusted to play back at a constant rate of 5 frames/s. Video corresponds to still images from Fig. 3 panel a.

**Video 4 – Figure 2 panel c – Laser ablation of a manipulated metaphase k-fiber under local microneedle load:** Laser ablation of an individual k-fiber after it was subject to microneedle manipulation force for 2 min without fracture reveals stability of the newly formed plus-end in representative PtK2 cell stably overexpressing GFP-tubulin. The microneedle (red, AF647 labelled, channel not shown after timepoint 0:00) is used to manipulate the k-fiber (white) before being ablated (yellow “X”) and its newly formed plus-end tracked (blue arrow), showing lack of depolymerization. Scale bar, 5 µm. Video was collected using a spinning disk confocal microscope at one frame every 10 s. Video has been adjusted to play back at a constant rate of 3 frames/s. Video corresponds to still images from Fig. 3 panel c.

**Video 5 – Figure 3 panel b – Bleach mark generation on a stabilized metaphase k-fiber stub after fracture was induced via microneedle load application:** Immediately after fracture via microneedle manipulation (microneedle channel not shown), a bleach mark was generated on the stabilized k-fiber stub (white) attached to the pole to monitor stabilized plus-end dynamics in representative PtK2 cell stably overexpressing GFP-tubulin. The bleach mark (magenta arrow) and subsequent tracking of the new plus-ends (blue arrow) reveals suppressed microtubule dynamics at the newly formed plus-end. Scale bar, 5 µm. Video was collected using a spinning disk confocal microscope (0 ± 0.6 µm, 0.3 µm z-spacing) every 10 s. A maximum intensity projection was generated at each timepoint to create the video. Video has been adjusted to play back at a constant rate of 2 frames/s. Video corresponds to still images from Fig. 1 panel j.

**Video 6 – Figure 4 panel a – Microneedle manipulation of an individual metaphase k-fiber in cell transiently overexpressing eGFP-doublecortin (DCX):** Microneedle manipulation of individual k-fiber reveals an example of DCX loss near the microneedle manipulation site in representative PtK2 cell stably overexpressing HaloTag-tubulin and transiently overexpressing eGFP-DCX. The microneedle (cyan, AF561 labelled, channel not shown after timepoint 0:00) exerts force on the k-fiber (magenta) and DCX (green) de-enrichment is seen. Scale bar, 5 µm. Video was collected using a spinning disk confocal microscope at one frame every 10 s. Video has been adjusted to play back at a constant rate of 3 frames/s. Video corresponds to still images from Fig. 4 panel a.

**Video 7 – Figure 5 panels a & b - Microneedle manipulation of an individual metaphase k-fiber in cell stably overexpressing GFP-EB1:** Microneedle manipulation of individual k-fiber reveals an example of fracture and GFP-EB1 enrichment at the stable newly formed plus-end in representative PtK1 cell stably overexpressing GFP-EB1 treated with SiR-tubulin. The microneedle (cyan, AF561 labelled, channel not shown after timepoint 0:00) exerts force on the k-fiber (magenta) and EB1(green) is enriched at the new plus-end (orange arrow). Following fracture a full cell volume was obtained (0 ± 4 µm, 0.3 µm z-spacing) and used to create a maximum intensity projection. Scale bar, 5 µm. Video was collected using a spinning disk confocal microscope at one frame every

10 s. Video was stopped once fracture occurred. Video has been adjusted to play back at a constant rate of 3 frames/s. Video corresponds to still images from Fig. 5 panel a.

**Video 8 – Figure 5 panels f & g - Microneedle manipulation of an individual metaphase k-fiber in cell stably overexpressing GFP-EB1:** Microneedle manipulation of individual k-fiber reveals an example of GFP-EB1 enrichment along the k-fiber prior to fracture that corresponds with the site of stabilization in representative PtK1 cell stably overexpressing GFP-EB1 treated with SiR-tubulin. The microneedle (cyan, AF561 labelled) exerts force on the k-fiber (magenta) and EB1(green) is enriched at a discrete site along the k-fiber (orange arrow). Scale bar, 5 µm. Video was collected using a spinning disk confocal microscope at one frame every 10 s. Video has been adjusted to play back at a constant rate of 3 frames/s. Video corresponds to still images from Fig. 5 panel a.

## Supplementary Figure Legends

**Supplementary Figure 1.**
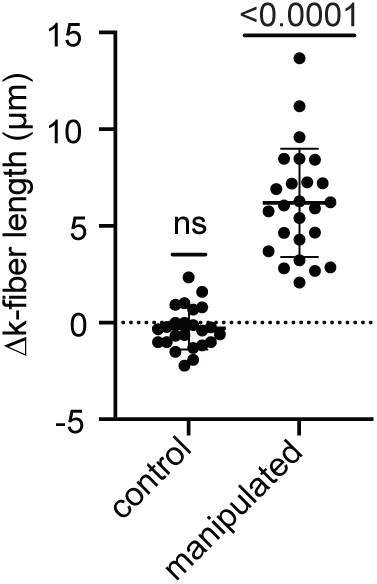
(corresponds to. Figure 1**): Microneedle manipulation causes k-fiber lengthening in only the manipulated k-fiber.** The differences in k-fiber length from the start of manipulation until the timepoint prior to fracture for manipulated k-fibers and a non-manipulated control k-fiber in the same cell. Change in length is measured as final k-fiber length – initial k-fiber length (one sample t-test, n=25 for manipulated, n=24 for non-manipulated, pooled across N=9 experimental days).

**Supplementary Figure 2.**
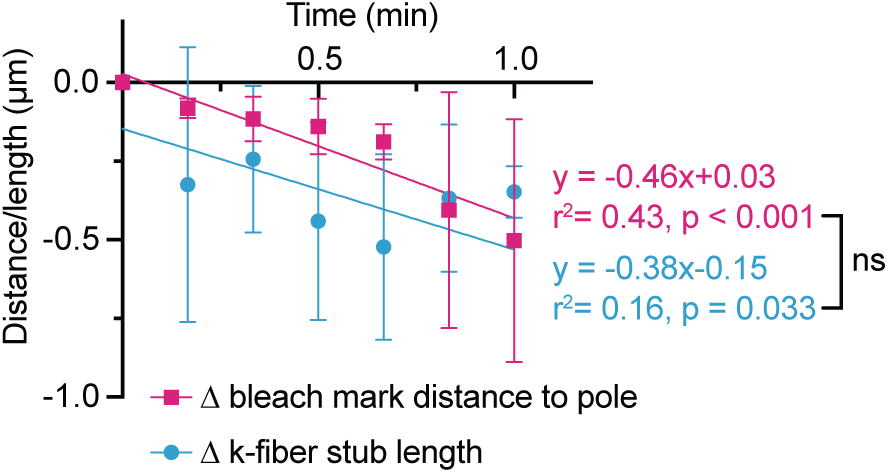
(corresponds to. Figure 3**): Newly formed plus-ends depolymerization speeds and k-fiber flux speeds are statistically indistinguishable post fracture.** The change in distance between the bleach mark and the pole and the newly formed plus-ends and the pole plotted over time (n=5 k-fibers, N=2 experimental days linear regression analysis, slopes tested for significant difference using a student t-test).

**Supplementary Figure 3.**
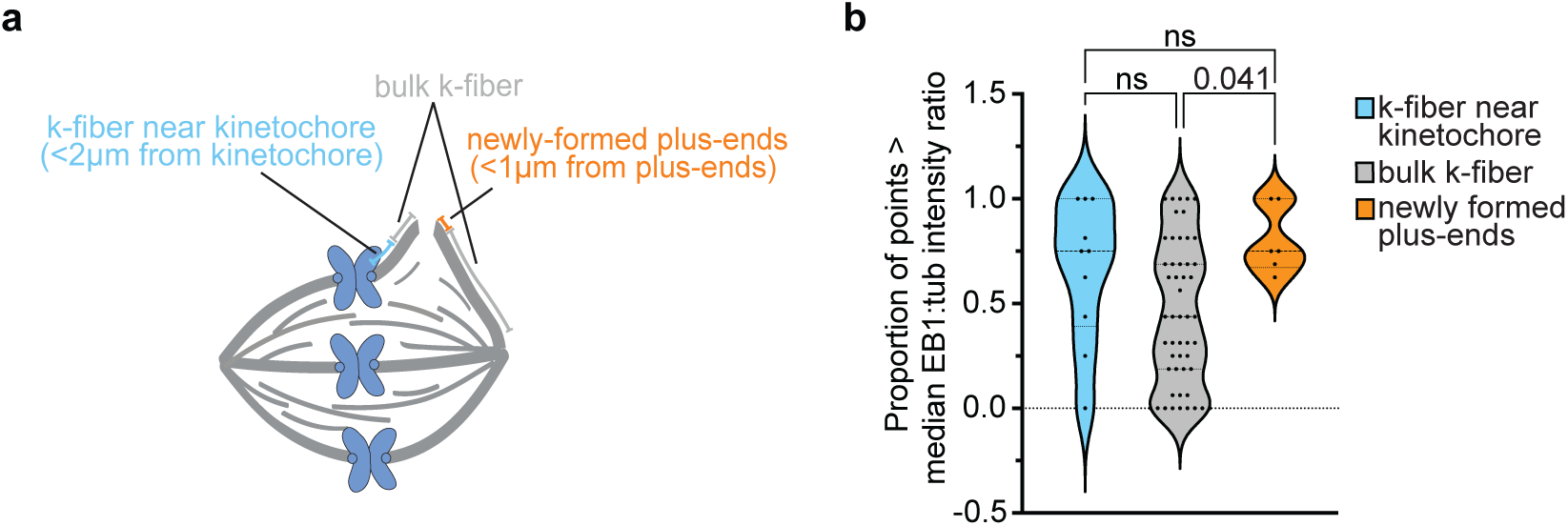
(corresponds to. Figure 5**): Newly formed plus-ends demonstrate local EB1 enrichment upon fracture. a)** Schematic depicting different regions within the manipulated spindle where the EB1:tubulin intensity ratio was quantified. **b)** Distribution of the percentage of points greater than the median EB1:tubulin intensity ratio along the manipulated k-fibers in PtK2 cells stably overexpressing HaloTag-tubulin and transiently overexpressing 2xGFP-EB1 binned by location (N=6 k-fibers, n=77 1-µm sections, M=3 experimental days, ordinary one-way ANOVA).

## Notes

### Competing Interest Statement

The authors have declared no competing interest.

